# Melanization is an important antifungal defense mechanism in *Galleria mellonella* hosts

**DOI:** 10.1101/2022.04.02.486843

**Authors:** Daniel F. Q. Smith, Quigly Dragotakes, Madhura Kulkarni, J. Marie Hardwick, Arturo Casadevall

## Abstract

A key component of insect immunity is melanin encapsulation of microbes. Melanization is also a part of an immune process known as nodulation, which occurs when insect hemocytes surround microbes and produce melanin. Insect nodules are analogous to mammalian immune granulomas. Melanin is believed to kill microbes through the production of toxic intermediates and oxidative damage. However, it is unclear to what extent immune melanin is directly fungicidal during infections of insect hosts. We reported previously that *C. neoformans* cells are encapsulated with host-derived melanin within hemocyte nodules. Here we report an association between melanin-based immune responses by *Galleria mellonella* wax moth larvae and fungal cell death of *C. neoformans* during infection. To monitor melanization *in situ*, we applied a tissue-clearing technique to *G. mellonella* larvae, revealing that nodulation occurs throughout the organism. Further, we developed a protocol for time-lapse microscopy of extracted hemolymph following exposure to fungal cells, which allowed us to visualize and quantify the kinetics of the melanin-based immune response. Using this technique, we found evidence that cryptococcal melanins and laccase enhance immune melanization in hemolymph. We used these techniques to also study the fungal pathogen *Candida albicans* infections of *G. mellonella*. We find that the yeast form of *C. albicans* was the primary targets of host melanization, while filamentous structures were melanin-evasive. Approximately 23% of melanin-encapsulated *C. albicans* yeast survive and break through the encapsulation. Overall, our results provide direct evidence that the melanization reaction functions as a direct antifungal mechanism in insect hosts.

## Introduction

Insects occupy essential niches in global ecosystems, including many that directly affect human health and survival ^1^. In addition, insects serve as powerful model systems for infectious disease research, and help to reduce reliance on vertebrates recommended by “3R” -Replace, Reduce, and Refine – programs ^2^. Insects are also targeted by environmental pathogens and have evolved complex immune mechanisms that partially overlap with mammalian innate immunity. Understanding the dynamics of insect-pathogen interactions and the factors involved is vital to both ensure ecosystem stability and establish invertebrate immunological models in research.

Fungi are an important class of pathogens for insects, and emerging fungal pathogens are predicted to become bigger threats to human health and agriculture in the coming years ^3,4^. Consequently, studying host-fungal interactions using insect models is important and timely. Although insects do not produce antibodies or other mammal-like adaptive immune responses, the antifungal immune defenses of insects involve cell-mediated and humoral innate immune processes ^5^. Hemocytes, the immune cells of invertebrates which circulate in the hemolymph, have roles comparable to macrophages and neutrophils in mammals. Hemocytes are responsible for clearance of fungi via phagocytosis, release of extracellular damaging reactive oxygen species (ROS) and inflammatory molecules, and the creation of granuloma-like structures through a process called nodulation ^6^. During nodulation, hemocytes surround the microbe and form an aggregate of insect cells, within which, clotting factors, immune enzymes, and immune complexes are released and activated ^6–8^. These structures immobilize the fungus and lead to its destruction. Also, during infection, the production of prostaglandins by the plasmatocyte subset of hemocytes in Lepidopteran species cause the lysis of other hemocytes called oenocytoids. The lysis of oenocytoid cells results in the release of antimicrobial peptides, signaling molecules, and enzymes important to immune function ^9–11^. One class of host enzymes that are often released and activated during oenocytoid lysis and nodulation are phenoloxidases (PO) ^9,11^. POs are enzymes responsible for converting catecholamines in the hemolymph into melanin ^12^. Melanin is a the black-brown pigment that is an important component of insect immune defense and wound repair ^13^. Melanization produces oxidative species and cytotoxic intermediates that are hypothesized to result in the death of the microbe ^12,14^. Additionally, melanin may act as a physical barrier, restricting gas exchange and nutrient uptake, and thus prevent fungal replication and dissemination to other tissues ^15^. At this time, *in vitro* evidence strongly links PO activity and resulting melanin intermediates with killing of fungi, bacteria, and viruses ^16–18^, but comparable direct evidence for the microbicidal effect of POs and their toxic intermediates *in vivo* during insect infections is challenging to measure directly. Consequently, obtaining direct evidence that the process of melanization is fungicidal *in vivo* is important for establishing insect melanin as an important mechanism for clearing fungal infections.

Larvae of the wax moth *Galleria mellonella* are commonly used as a model organism for studying fungal pathogenesis ^5^. *G. mellonella* larvae are readily available in large numbers at low cost. Their larger size (2-3 cm) relative to other model insects such as *Drosophila melanogaster* makes them amenable to research approaches requiring larger volumes of hemolymph, insect hemocytes, and soluble immune factors. The study of *G. mellonella* hemolymph can prove valuable for understanding the insect’s immune response to infection and stress. *G. mellonella* are also commonly used as a model for studying mammalian pathogens, including human pathogenic fungi *Cryptococcus neoformans* and *Candida albicans* ^19–21^. While *G. mellonella* is a model for mammalian fungal infections because of similarities between the *G. mellonella* immune responses and the mammalian innate immune responses ^5^, a more thorough understanding of the insect immune response is needed to fully benefit from studying host-microbe interactions in *G. mellonella*.

The differences between mammalian versus insect hosts also provide important new insights into host-microbe interactions and mechanisms of fungal virulence factors ^5,19,20,22,23^. For example, laccase, a fungal enzyme that oxidizes mammalian and insect catecholamines, is an important virulence factor in both hosts but, seemingly by distinct and diverse mechanisms ^19,24,25^. In insects, fungal laccase appears to oxidize and deplete host catecholamines required for encapsulating the fungus in melanin, thus weakening the host immune response. Fungal laccases also help detoxify reactive oxygen species that form during insect immune processes ^24^. In contrast, during mammalian infection, fungal laccase enhances production of fungal melanin to evade key mammalian immune defenses ^26^. Thus, fungal melanins increase virulence in mammals, but decrease virulence in *G. mellonella* ^27–29^. These seemingly different and opposite roles in which fungal melanins interact with mammalian and insect hosts is unexplored and as of now unexplained in literature.

In this study, we describe the first direct evidence that the melanin-based immune response *in vivo* is fungicidal against *C. neoformans*. Our data show a direct link between melanin encapsulation during infection within the *G. mellonella* larvae and fungal death by visualizing death using an endogenously expressed GFP-based fungal viability assay. We then used a series of *in situ, in vivo*, and *in vitro* methods to study the melanin-based immune response of *G. mellonella* larvae. For *in situ* experiments, we modified a previously published tissue-clearing protocol to visualize melanized nodules and their tissue specificity, or lack thereof. We have also developed time-lapse microscopy method for visualizing the melanin-based immune system. We applied this method to quantify the melanization kinetics during *in vitro* fungal infection, which improved our understanding of how fungal components, such as laccase and melanin, interact with insect melanization. We gained insight into how *Candida albicans* activates and evades the melanin-based immune response through morphological switching. Overall, our findings strongly suggest that melanization has direct antimicrobial activity *in vivo* in the insect immune system, and we subsequently explore methods to further study the melanization immune response.

## Results

### *Galleria mellonella* kill *C. neoformans* through melanin encapsulation in nodules

Previously, we found that *C. neoformans* is encapsulated inside immune system-produced melanins during infection of *G. mellonella*, providing evidence that the melanin-based immune response is activated against *C. neoformans* in *G. mellonella* ^30^ (Figure 1A). To evaluate whether insect melanin encapsulation kills *C. neoformans*, we assessed viability using a GFP-expressing strain of *C. neoformans*, which expresses GFP under an actin promotor. The GFP-expressing strain as a reporter for fungal viability *C. neoformans* was validated using the standard dead cell stain propidium iodide. Propidium iodide staining was nearly mutually exclusive with GFP fluorescence in untreated cells, and GFP fluorescence was extinguished when heat killed (Figure 1B, Supplementary Figure S1A and S1B). Using GFP fluorescence as a proxy for cell viability, we found fewer GFP-positive fungal cells in association with nodules located in the hemolymph compared to non-melanin encapsulated fungal cells at both room temperature and at 30°C at 24 and 72 h post-infection (Figure 1C and D). Melanin produced by *C. neoformans* in culture did not quench or obscure the GFP fluorescence, as determined by imaging melanized versus non-melanized *C. neoformans* H99-GFP (Supplementary Figure S1C-D). Hence, this result suggests that the immune melanization reaction itself is associated with fewer GFP-positive cells, consistent with death of *C. neoformans in vivo* during infections of *G. mellonella*.

**Figure 1.**
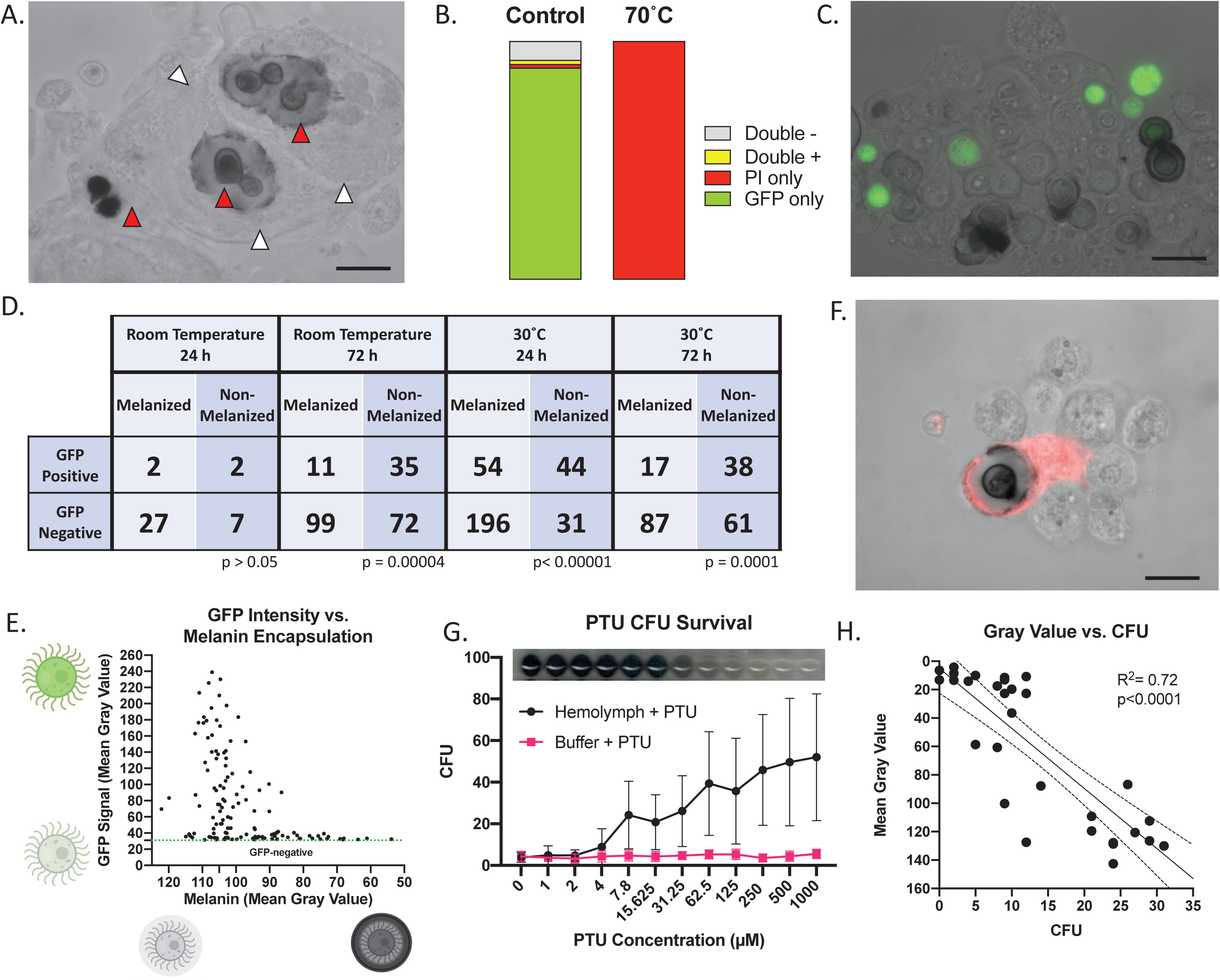
The melanin-based immune response is implicated in killing C. neoformans during infection. (A) During infection of *G. mellonella*, melanized nodules are formed within the hemolymph, which encapsulate the fungus. Red arrows indicate melanin-encapsulated fungus, while white arrows show the hemocytes surrounding the fungi. (B) GFP fluorescence is lost in heat killed *C. neoformans* cells expressing GFP under an actin promoter. (C) Using a GFP-expressing strain of *C. neoformans*, we can visualize cell viability within the nodule, with living fungi fluorescing green and dead cells having no signal. (D) Melanin-encapsulated fungi are statistically less likely to have a GFP-positive signal at 24 h after infection at 30°C, and at 72 h post infection at room temperature and 30°C (p<0.00001, p=0.00004, and p=0.0001, respectively, Chi-squared test), indicating that melanin-encapsulation is associated with fungal death. (E) In GFP-positive cells, the brighter fluorescence signal correlates with little to no melanization, with no cells that have strong GFP signals and large amounts of melanin encapsulation. Each point represents data from one GFP-positive fungus within a nodules from three biological replicates. (F) Propidium Iodide viability staining does not appear to penetrate the inner space of the nodule where the fungus is located, but the staining does show that the hemocytes surrounding the fungus are not viable. (G) Colony forming units (CFUs) correlate directly to melanin inhibition using PTU (inset) in vitro, which indicates that melanin has a role in controlling growth of C. neoformans. Each data point represents the mean and 95% Confidence Interval of three independent biological replicates. (H) CFUs also inversely correlated to the Mean Gray Value of these wells, meaning that the darker wells with more melanization had higher CFUs recovered compared to the melanin-inhibited wells. Each data point indicates measurements from individual wells across three biological replicates. (A-C,F) Data shown is representative of three biological replicates. All experiments performed in biological triplicate, with (G) representing 5 biological replicates. Scale bars represent 10 µm.

Within nodule-encapsulated GFP-positive *C. neoformans*, we measured the degree of immune melanin intensity and GFP fluorescence intensity. We found that the yeast with the weakest GFP signal tended to be encapsulated with more melanin surrounding them, compared to the population of brightly fluorescent cells (Figure 1E). The result was an inverse correlation between melanization and fluorescence within GFP-positive cells. Most GFP-positive cells had little to no melanization, with a mean gray value around 105, which is the background gray value intensity. The occurrence of faint signal in some cells suggests that these emanate from cells in the process of dying or having been recently killed.

We attempted to use PI as an additional technique to study fungal viability within the nodules and to show that the GFP-negative cells are indeed dead. Surprisingly, PI staining did not result in the expected fluorescence in nodule-encapsulated yeast cells, but there was staining in some of the hemocytes that surrounded the yeast cells, and the external periphery of the nodules (Figure 1F). Given that PI staining was extracellular to the fungal cell, this staining could reflect released fungal or hemocyte DNA. The absence of PI staining for fungal cells in nodules suggests that PI was unable to reach the center of the nodules where the fungi are found and shows that some of the hemocytes involved in surrounding the fungus may undergo cell death in the process. (Figure 1F). The permeability or access issues that may arise when using added dyes to measure microbial viability within the nodule show the usefulness of using a live-dead indicator that is endogenously present within the fungus, such as the constitutively expressed GFP.

To confirm that insect melanin killed *C. neoformans*, we performed complementary experiments *in vitro* by assessing the ability *G. mellonella* melanization to inhibit the growth of *C. neoformans*. We incubated *C. neoformans* cells with extracted hemolymph from *G. mellonella* in a 96-well plate. Using various concentrations of a phenoloxidase-specific competitive inhibitor, phenylthiourea (PTU), we were able to generate a range of melanin inhibition conditions in the hemolymph-fungal mixture. After 24 h, we removed a small aliquot of the mixture and plated it on nutrient rich agar to allow fungal growth. The number of CFUs following a 24 h incubation with the hemocytes and PTU melanin inhibitor was directly proportional to the concentration of PTU (Figure 1G), and thus inversely proportional to the degree of melanization (Figure 1G, inset). Melanization, as measured by mean gray value, correlated with low CFUs (Figure 1H). This result strongly suggests that immune melanization, inhibits the growth of *C. neoformans in vitro*.

### Time-lapse microscopy of hemocyte-fungal interactions and the melanization response

To record the kinetics of the hemolymph melanization response, we developed a protocol to extract hemocytes and watch their interactions with fungi (Figure 2A). Using this time-lapse microscopy, we were able to measure the rate and magnitude of the anti-cryptococcal immune melanization response following treatment with different species of fungi, mutants, or isolated virulence factors (Supplementary Video 1-2). Using particle analysis, we could quantify the area covered by melanization in minute intervals, and we can record the rate of hemolymph melanization (Figure 2B-D). By analyzing the melanization kinetics between different mutant and wildtype strains, we can determine how the mutant gene of interest affects fungal interactions with the melanization immune response. We observed variation in the extent of melanization between different experiments, likely due to variability in the fungi in the field of view, and biological variability from non-isogenic *G. mellonella* larvae. To overcome the risk of interpreting variability-derived artifacts as results, each experiment was performed with a corresponding control (i.e., parental strain and mutant experiments were performed at same time, using the same stock of hemolymph and the same pool of extracted hemocytes).

**Figure 2.**
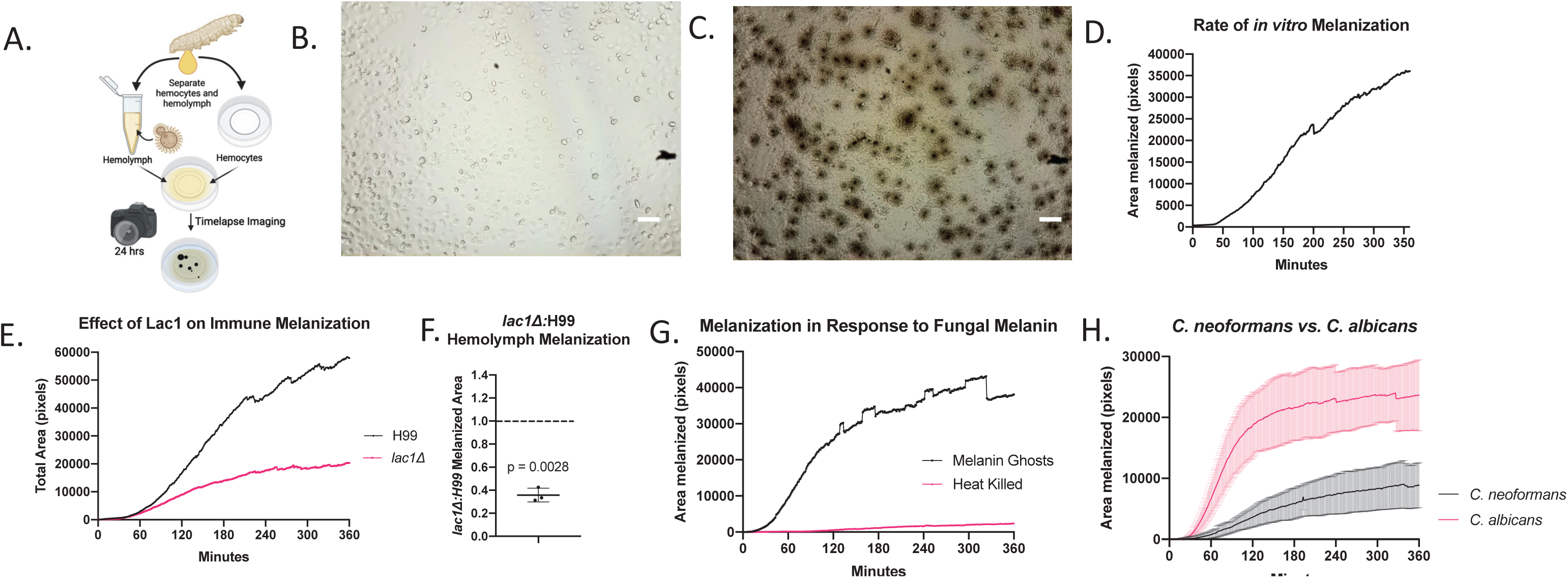
*Using* in vitro *timelapse microscopy to visualize the melanin-based immune response*. (A) Using a developed timelapse microscopy protocol, we were able to record the dynamics of hemolymph melanization in response to *C. neoformans* from 0 (B) up to 24 hours (C). We can then use particle measurement software to visualize and quantify the melanization reactions over time (D). This method can be used to gain novel insight into how different virulence factors, such as laccase, influence the melanization response, where we see that the laccase knockout mutant causes less of a melanization response without much change in time until onset of melanization (E). Overall, the *lac1Δ* triggers ∼40% as much of a melanization immune response compared to the WT (F). (G) Isolated fungal melanins are associated with activation of the insect melanin based immune response, whereas the heat killed cryptococcal cells are not (H) Additionally, we can compare the activation of the melanin-based immune response from different fungal species such as *C. neoformans* and *C. albicans*, which strongly activates the melanization immune response. (B-D, G) are representative images and graphs from three biological replicates. (E,F,H) represent averages from biological replicates. Error bars indicate Standard Deviation. Scale bars represent 50 µm.

When we evaluated the rate and magnitude of hemolymph melanization in response to the *lac1Δ* mutant, we found that there was a dramatically reduced rate and magnitude of hemolymph melanization compared to the wildtype parental strain (Figure 2E). While the overall magnitude of melanization varied between replicates, the ratio of insect melanization that occurred in the *lac1Δ* versus H99 (WT) was statistically significantly lower than 1, and consistently around 0.35 (Figure 2F).

Many fungi, including most of those that infect people, produce melanin in their cell walls, which enables them to persist within mammalian hosts and avoid destruction from oxidative stress and antimicrobial agents ^26^. However, literature shows that melanized fungi are less virulent in *G. mellonella* compared to their non-melanized or albino mutant counterparts ^27,28,31^. We hypothesized that the fungal melanin could act as a pathogen or damage-associated molecular pattern (PAMP/DAMP), resulting in enhanced activation of the melanization immune reaction. We found that isolated *C. neoformans* melanin activated immune melanization, both with (Figure 2G) and without hemocytes present compared to heat killed *C. neoformans* (Supplementary Video 3-5). Since melanin ghosts contain trace amounts of fungal cell wall components that could theoretically activate the melanization immune response, we used heat-killed non-melanized cells as a control for the cell wall components that would be present. This indicates that there is a mechanism by which fungal melanin is specifically recognized and activates the phenoloxidase cascade. Further, the isolated melanin ghosts are aggregated by the hemocytes throughout the course of the time-lapse microscopy, even when immune melanization is not activated (Supplementary video 6).

We used this technique to compare the immune melanization between *C. neoformans* and *C. albicans*, the latter of which is known to trigger robust melanization of the hemolymph. In the time-lapse microscopy, we saw that *C. albicans* activated the melanization response faster (beginning as early as 15 minutes) and to a significantly greater extent than did *C. neoformans* (Figure 2H). This corresponds to the levels of melanization previously reported that occurs during *G. mellonella* infection with *C. albicans* versus *C. neoformans* and validates that our system, at least in part, corresponds to what occurs during actual infection.

### Evaluating the melanin-based immune response of *G. mellonella* using tissue clearing

Tissue clearing is a technique that allows for visualization of structures deep within an organism or tissue sample, without significant disruption of the native tissue anatomy. We adapted a previously reported protocol ^32^ to visualize the anatomical localization of the anti-cryptococcal melanization response in *G. mellonella* (Figure 3A,B). We found that using this technique, we could visualize melanized nodules *in situ* that are formed only during infection with *C. neoformans* and not in uninfected controls (Figure 3C,D). These *in situ* melanized nodules (Figure 3D-F) appeared very similar to those that are collected from extracted hemolymph, which represent an *in vivo* method of visualizing the nodules (Figure 1A,3G). The visual similarities between Figure 3E,F and Figure 3G clarified that the nodules observed in extracted hemolymph are generally representative of the entirety of nodules in the organism. Both the *in vivo* and *in situ* techniques could be quantified to determine the average melanized nodule area and degree of melanization (Figure 3H). However, the cleared tissue had some opacities or normally darker tissues (i.e. digestive tract contents, legs, prolegs, spiracles, cuticle pigmentation, etc.), which can result in the detection of dark particles even in the uninfected controls, albeit at a much lower frequency (Figure 3H). Further, while there are some large *C. neoformans* nodules *in situ* that appeared aggregated together (Figure 3I, arrows), there was no clear anatomical tropism for nodule formation and the nodules are found throughout the larvae, implying that the infection was disseminated throughout the body of larvae, possibly through the insect’s open circulatory system. The large, aggregated nodules can be imaged along the Z-axis, which allowed 3D reconstruction of the nodule for a better understanding of the native nodule structure compared to the *in vivo* preparations compressed under a slide (Supplementary Video 7). However, compared to the *in vivo* experiments, the resolution of the melanin-encapsulated *C. neoformans in situ* is limited, and variations in opacity and tissue thickness could interfere with measurements.

**Figure 3.**
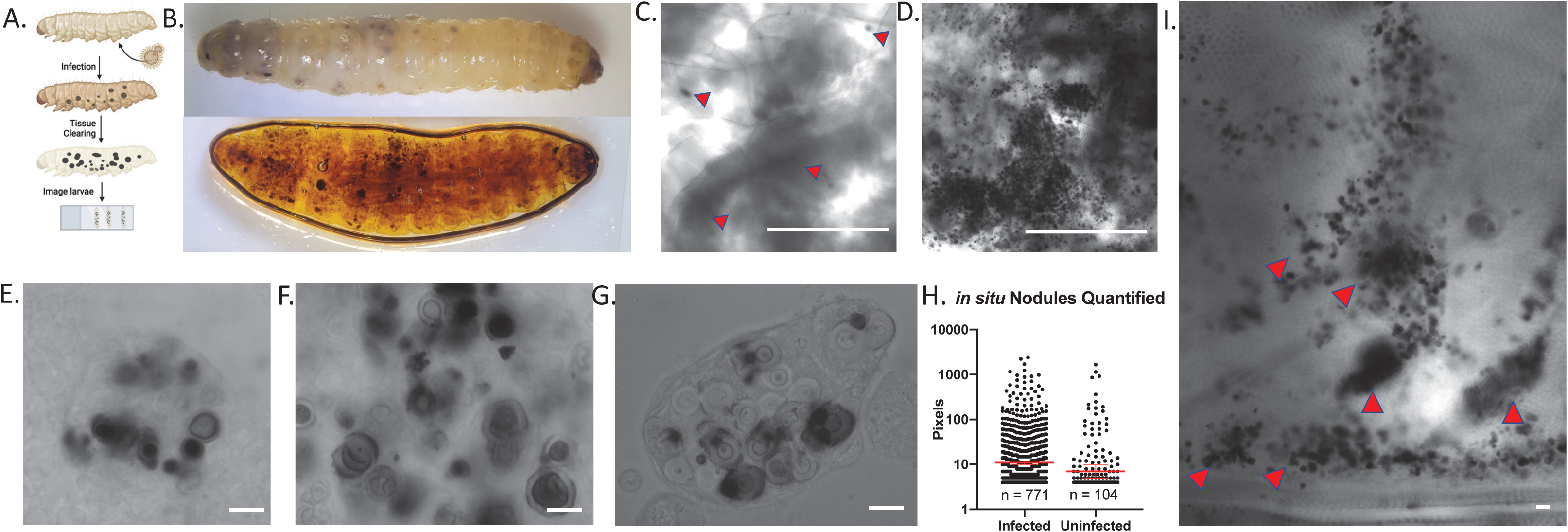
*Using tissue clearing to visualize the melanin-based immune response against* C. neoformans in situ. Using tissue clearing techniques (A), we allowed for better visualization of melanized nodules within the intact *G. mellonella*, as seen in a before (top) and after (bottom) view of the same infected larvae (B). Under microscopy, the uninfected larvae have little to no melanized nodules (red arrows indicate dark, irregular, or opaque areas in the tissue) (C), whereas the *C. neoformans* larvae have very clear and distinct nodules (D). These nodules can be viewed under high magnification, where the nodule structure and encapsulated fungus is apparent (E-F). These structures are very similar in appearance to the nodules extracted from hemolymph (G). The size of the melanized nodules can be quantified using particle measuring software. Infected particles represents n = 771, uninfected particles represent n =104, p = 0.016 (Mann-Whitney test) (H). (I) The melanized nodules of *C. neoformans* appear to have no particular tissue tropism, but can be found in distinct clumps or areas throughout the larvae (red arrows). (B-G, I) are representative images from three biological replicates. Scale bars in (E-G, I) represent 10 µm, and in (C,D) represent 500 µm.

### *Candida albicans* with the Melanin-based Immune Response

*C. albicans* is a fungus known to elicit a strong melanization reaction in hemolymph of infected *G. mellonella* larvae^33^. We thus employed the *in vitro, in vivo*, and *in situ* techniques described above to gain insight into the host-microbe interactions of *C. albicans* with the *G. mellonella* melanin-based immune response.

Using the *in vivo* technique of extracting infected hemolymph to analyze melanized nodules, we observed melanin-encapsulated *C. albicans* cells within nodule structures (Figure 4A). These melanized nodules are like those observed during *G. mellonella’*s infection with *C. neoformans*, however, the borders of the melanin itself appeared less distinct, blurry, and smudged. An additional difference from cryptococcal infection was the presence of filamentous *C. albicans* structures within the nodules. These hyphae or pseudohyphae were melanin-encapsulated, but seemingly to a lesser extent than the yeast morphology.

**Figure 4.**
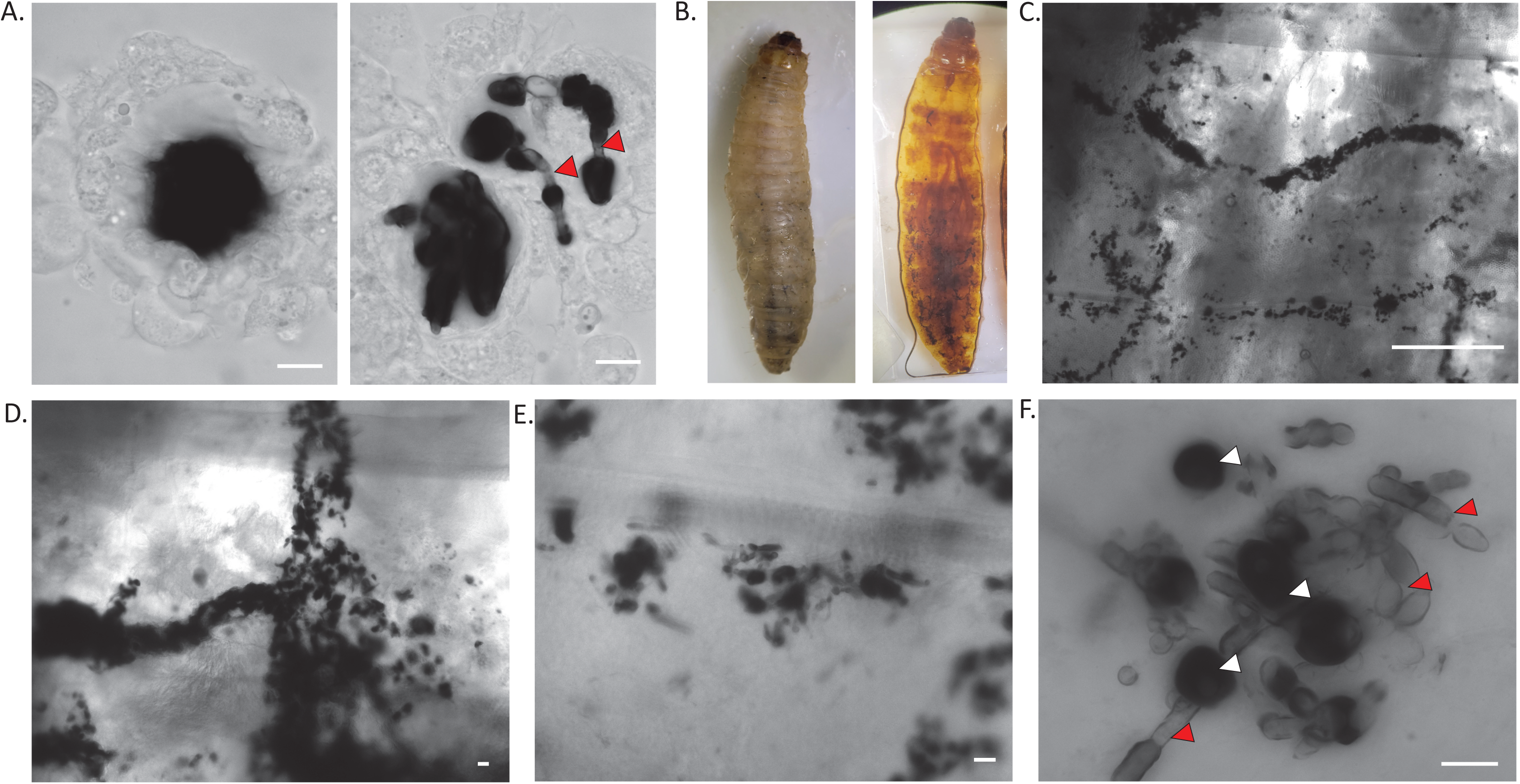
Visualizing the melanization response to infection with Candida albicans. (A) Similar to the hemolymph from larvae infected with *C. neoformans*, we are able to see melanized nodules within the hemolymph. The melanized spot within these nodules appears more diffuse/less defined, and in some, the presence of less-melanized hyphae is distinct (red arrows). (B) We can also use tissue clearing to visualize the melanized nodules during *C. albicans* infection. (C-D) The melanized *C. albicans* seem to cluster in specific areas, in long strips within the larvae. (E-F) Under higher magnification, we can see hyphal structures of *C. albicans*, which corresponds to what is previously known about *C. albicans* morphology in *G. mellonella*. Interestingly, the hyphae appear less melanized (red arrows) compared to the spherical yeast (white arrows) (F). All panels show representative images from 3 biological replicates. Error bars in (A,D-F) represent 10 µm and in (C) represent 500 µm.

Analysis of the tissue of larvae infected with *C. albicans* for 24 h using the *in situ* tissue clarification technique (Figure 4B) revealed groupings of the melanized nodules, often in long string-like patterns (Figure 4B), that did not appear particularly associated with any organs or tissues (Figure 4C-D, supplemental Figure 2A-F). Under higher magnification, we observed lightly pigmented hyphae and dark spherical melanized particles within the cleared larvae after 24 h of infection (Figure 4C-F). These images validate that filamentation occurred within *G. mellonella*, which was previously observed with histology. The melanin-encapsulated fungi form large aggregates (Figure 4C,D), which we initially thought could be indicative of a tropism for a specific tissue such as the chitinous trachea. However, upon dissection of uncleared infected larvae, there did not appear to be an association of these clusters with any specific tissues (Supplementary Figure 2 A-F). Differences in pigmentation between the two *C. albicans* morphologies, particularly as seen in the Z-projection in (Figure 4F), indicated that the hyphae were encapsulated with less melanin during infection compared with the yeast form of the fungus. One potential bias in interpreting this data is that since we are only looking at melanin pigmentation, we are likely missing any non-melanin encapsulated fungi which would blend in with the insect tissue.

We used the *in vitro* time-lapse microscopy to observe the melanization dynamics of *C. albicans* in hemolymph. As seen earlier (Figure 2H), *C. albicans* triggered a more robust melanization response than *C. neoformans* (Figure 5A). In time-lapse microscopy performed without the addition of insect hemocytes, we observed that the *C. albicans* began to grow in filamentous forms (Figure 5B2), consistent with the importance of filamentation in the pathogenesis of *C. albicans* within *G. mellonella* and the *in situ* data (Figure 3)^23^. In mammalian hosts, filamentation is triggered by serum, neutral pH, and temperature ^34,35^. However, in the *G. mellonella* system, filamentation *in vitro* does not occur when the *C. albicans* is only incubated with hemocytes without hemolymph, indicating that a component of the hemolymph is necessary for the morphological switch. Interestingly, as the time-lapse movie progressed, we observed that the hyphae did not get encapsulated by melanin in comparison to the yeast form of *C. albicans* (Figure 5B2). In mammalian hosts, filamentation by *C. albicans* is used to evade immune detection in part due to changes in cell wall structure and expression that prevent binding of C-type Lectins. After about 12 hours of filamentous growth, we observed the formation of blastoconidium (yeast) along the hyphae (Figure 5B3). The formation of these yeast cells then corresponded with a subsequent “bloom” of melanization (Figure 5A, B4, Supplementary Video 8). A similar temporal progression of *C. albicans* morphology and melanin-encapsulation is seen in *C. albicans* infected larvae dissected at various timepoints post-infection (Supplementary Figure 2G). The average time of this melanin bloom was about 840 minutes, with a 95% confidence interval between approximately 720 minutes and 960 minutes (Figure 5C).

**Figure 5.**
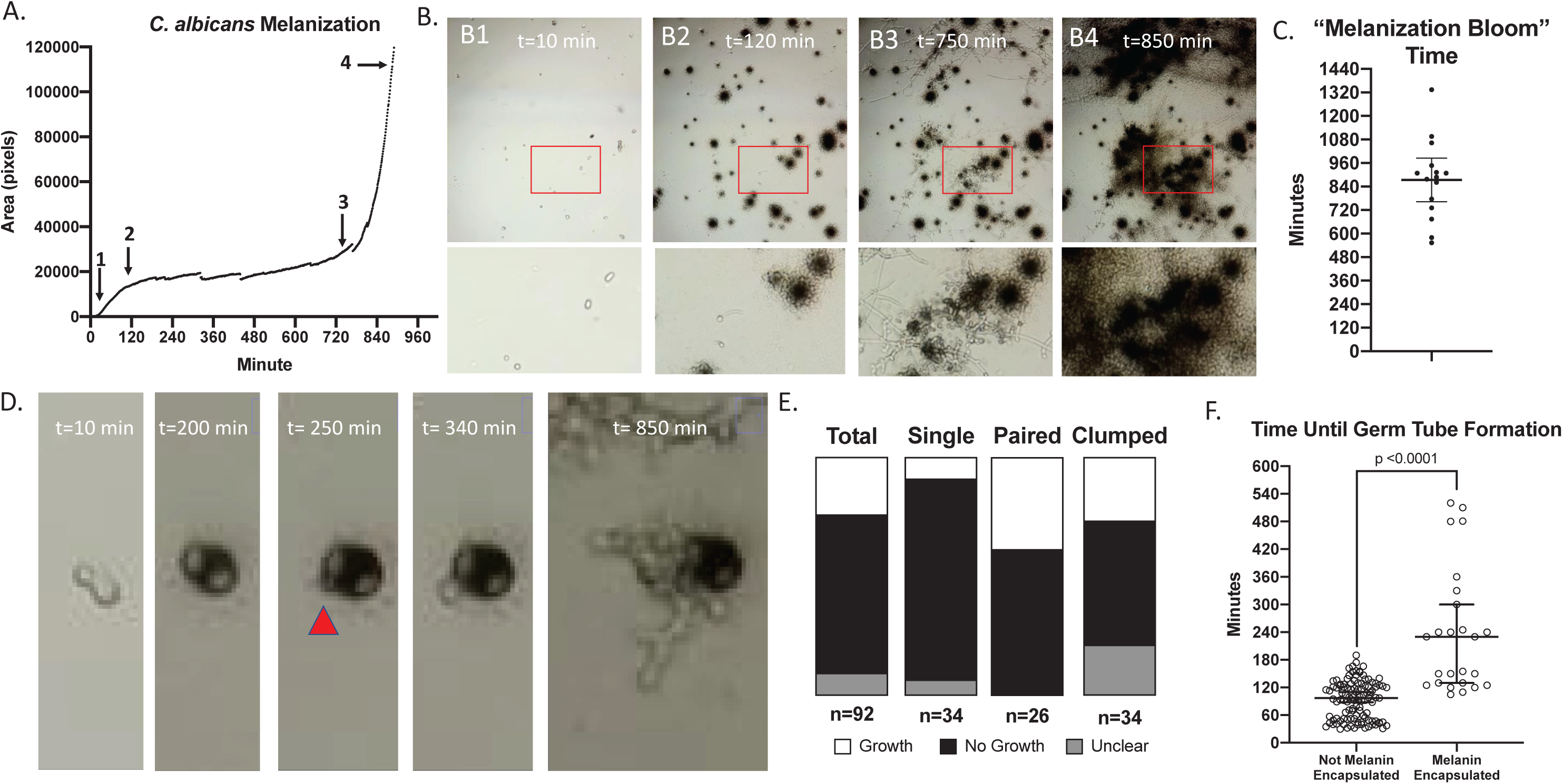
*Using timelapse microscopy to gain insight into* C. albicans *yeast and hyphae interactions with the melanization immune response*. (A) Kinetics of the melanization reaction within hemolymph incubated with *C. albicans* yeast, with a distinct tri-phastic response in which there is an initial peak reaction, an extended plateau, and a greater second peak of the melanization reaction. (B) Using microscopy, we can see that the yeast start off non-melanized (B1), and most reach their peak melanin-encapsulation by 120 minutes (B2). After which, the yeast begin to filament, and hyphae grow between 120 min and 750 minutes with minimal melanin encapsulation occurring. At 750 minutes (B3), blastoconidium (yeast) begin to form on the hyphae, which then corresponds to a rapid increase in melanization (B4). (C) The time until this melanin bloom is approximately 14 hours. Through these movies, we also see that some melanin-encapsulated yeast are able to survive the immune response and grow from underneath the layer of pigment (D). The melanin-encapsulated *C. albicans* in panel (D) begins to grow through the melanin by 250 minutes. (E) Overall, approximately 23% of melanin-encapsulated yeast are able to grow following melanin-encapsulation, with pairs of melanin-encapsulated yeast having the highest percentage of growing, with a 33% occurrence. (F) In the yeast that are able to grow following melanin-encapsulation, the time until germ tube formation is delayed to on average 230 minutes (n = 26), while in non-encapsulated yeast, the average time is 90 minutes (n = 111). Statistical test represented in (E) is an unpaired t-test, and error bars indicate Median with 95% Confidence Interval. (A,B,D) show representative data from three biological replicates. Each data point in (C) represents an independent replicate. (E,F) show data from 4 biological replicates. Error bars indicate 100 µm.

We observed that some of the melanin-encapsulated yeast survived the immune reaction and then underwent hyphal and/or pseudohyphal growth (Figure 5D, Supplementary Video 9). This occurred in about 23% of melanin-encapsulated yeast, with 8% of single yeast and 38% of budded/pairs of yeast being able to escape (Figure 5E) The time until hyphal or pseudohyphal growth was significantly delayed in the melanin-encapsulated cells; the median time for a non-melanin encapsulated *C. albicans* cells to begin filamentation is 97 minutes, while the melanin-encapsulated counterparts take 230 minutes, with some taking as long as 520 minutes (Figure 5F). This delay could be reflective of physical barriers as the fungus breaks through the melanin layer and/or delays in initiating cellular growth because of cell damage caused by the immune response.

Altogether, these data demonstrate three phases of the *C. albicans-*immune melanization interactions: 1) yeast become encapsulated with melanin, with nearly 25% surviving and breaking through the pigment, 2) cells undergo a yeast-to-hyphal transition, with the hyphal and pseudohyphal cells evading the melanization immune response; and 3) filamentous *C. albicans* begins to produce more yeast cells (referred to as blastoconidia or blastospores), which then causes a second bloom of melanization to occur (Summarized in Figure 6). While host melanization and fungal filamentation have been well-reported during the course of *C. albicans* infection^23,36–39^, this is the first indication that the hyphae and pseudohyphae are melanin-evasive. Although the presence of blastoconidia has been reported in *G. mellonella* infected with *C. albicans* ^36^, these data indicate for the first time that lateral blastoconidia growing from hyphae induce a strong “second wave” melanization response.

**Figure 6.**
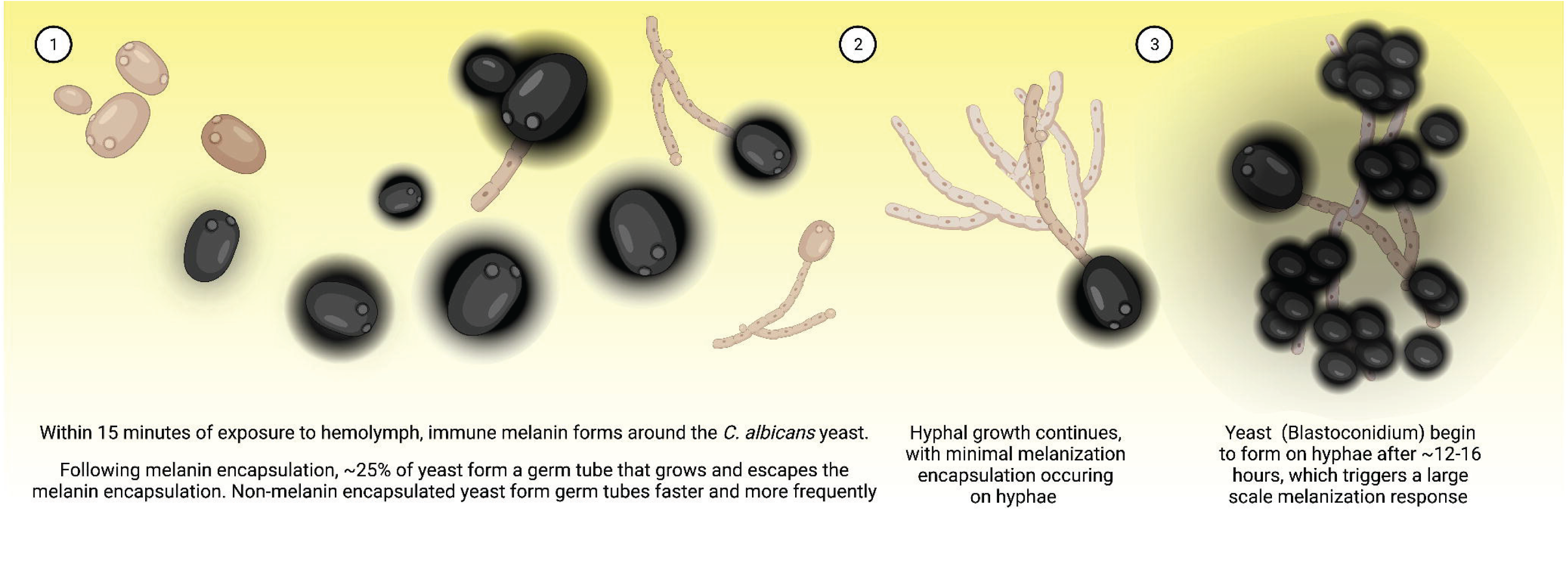
Overview of the growth of *C. albicans* within *G. mellonella* and the fungus’ interaction with the melanization immune response.

## Discussion

Melanin has been appreciated as a key part of the insect immune defense against microbes and parasites for the greater part of the past century ^13,40^. Immune melanization has been implicated as a major process in neutralizing entomopathogenic fungi upon infection^41^. The insect phenoloxidase and the melanization cascade produce toxic intermediates such as dihydroxyindole (DHI) and high levels of oxidative stress that can overwhelm and kill the fungus or microbe in vitro ^13,17^. However, there have been no *in vivo* studies showing that melanization directly kills fungi during these immune reactions within the insect. In this paper, we fill that gap by showing that melanization within nodules is associated with the death of *C. neoformans* using a GFP viability reporter assay and provide additional *in vitro* data for a fungicidal role in immune defense.

During cryptococcal infection of *G. mellonella*, melanin encapsulation of the fungus melanin encapsulation within nodules was associated with diminished or lost fluorescence signal in these GFP-expressing *C. neoformans* strains. Additionally, the melanin-encapsulated fungi that remained GFP positive had weaker signals and the intensity of the GFP signal was more intense for the non-melanin encapsulated fungi within the nodules. The expression of GFP in these cells is under the control of an actin promotor, and while actin is generally presumed to be constitutively expressed in cells, growth conditions have been shown to lead to some alterations in cryptococcal actin expression ^42,43^. If the environmental conditions within the nodule abolished actin expression in some cells without killing the fungus, we would expect that condition to equally affect the melanin and non-melanin encapsulated fungi, and as a result, see similar GFP-negative: GFP-positive ratios between the melanin-encapsulated and not melanin-encapsulated cells. The association between melanin encapsulation and disappearance in GFP fluorescence provides strong evidence for the notion that the melanization reaction kills fungal cells during infection. This is the first evidence that *G. mellonella* immune melanization directly and effectively neutralizes *C. neoformans* during infection and the first demonstration that melanin encapsulation results in fungal death within the insect. Previously, the death of microbes, specifically bacteria, was attributed to the enzymatic activity of the melanin-producing phenoloxidase (PO) in an *in vitro* reaction ^16^. In addition to our association of melanin encapsulation and fungal death *in vivo*, we sought to reproduce these results *in vitro* using extracted hemolymph in buffer. We used the PO-specific inhibitor, phenylthiourea (PTU), to inhibit PO activity and melanization and found that PO-inhibited wells of hemolymph had higher recoverable CFUs of *C. neoformans* compared. The inverse correlation of melanization with CFUs further supports the claim that melanin plays a role in neutralizing *C. neoformans*. Since we only assayed CFUs from these *in vitro* experiments, we cannot determine whether the melanization in the *in vitro* experiments directly killed the fungus or just inhibited fungal growth.

In addition to studying the extracted hemolymph, we used a developed *in vitro* time-lapse microscopy assay. We investigated the impact of fungal melanins on the insect immune response. We found that isolated fungal melanins, termed “melanin ghosts,” activated the melanin-based immune response whereas the heat-killed *C. neoformans* did not. This suggests that fungal melanins can activate the immune system which could help promote fungal clearance. This is interesting in the context of naturally-occurring fungal pathogens of insects, which tend to have a white color and/or do not naturally produce melanin pigment, such as *Beauveria bassiana* and *Metarhizium anisophilae* ^24,25^. This may be because of evolutionary pressure that selects for entomopathogenic fungi that produce less fungal melanin and thus are better at evading the insect’s melanin-based immune response. Since melanin is a component of the insect wound response, it is possible that these exogenous melanins are recognized by the insect as a damage associated molecular pattern (DAMP) and launches an inappropriate wound repair response.

We found that the *lac1Δ* mutant, which is unable to produce the enzyme laccase, causes less melanization in the hemolymph. This implies that some of the trigger for melanization comes from laccase-catalyzed initiation of melanin formation using host-derived catecholamines in the hemolymph. This is consistent with the observation that for *B. bassiana* infection of *G. mellonella*, that laccases play a role in virulence by oxidizing the hemolymph catecholamines and preventing them from producing anti-fungal melanization and reducing the oxidative burden on the fungus ^24^. It is also worth noting that the *lac1Δ* mutant is less virulent in *G. mellonella* infections compared to the parental ^19^. Together, these observations paint a nuanced picture of the role that laccase and fungal melanin play during fungal pathogenesis in *G. mellonella –* both fungal melanin and fungal laccases activate the melanin-based immune response, while fungal melanins are associated with decreased virulence, fungal laccases enhance virulence. We note that laccase is secreted by *C. neoformans* and is found in extracellular vesicles, which could transport laccase away from the fungal cell and reduce the antifungal damage from its effects on triggering insect immune melanization. We were also able to compare the amount of melanization that *C. neoformans* triggers with the amount triggered by other fungal species such as *C. albicans*. The differences in hemolymph-induced melanization during exposure to *C. albicans* and *C. neoformans* were previously described ^33^, and our results confirm those findings.

The second method used to evaluate the melanization response to *C. neoformans* was tissue clarification, which enabled us to visualize melanized nodules *in situ* deep within the larvae. We modified a recently developed protocol ^32^ to view the nodules that formed during infections, and saw the native structures of the nodules and their anatomical location in the larvae. This offers an advantage over dissection of uncleared larvae, because during the dissection process: 1) the tissue organization is disrupted, 2) some organs such as the nerve cord and cardiac system might be disrupted, and 3) the geography of infection patterns may not be apparent. Additionally, melanized nodules may not be visible within or behind opaque tissues and organs. Tissue dissection of opaque larvae was helpful when evaluating tissue tropism since tissue boundaries may not be fully visible in clarified larvae. A bias involved in studying fungal infections using both tissue clearing and dissection is that the non-melanized nodules or fungi may be missed, as unpigmented fungi will likely blend in with surrounding tissue. However, in the clearing method, we viewed the nodules throughout the entire depth of the larvae at a low to moderate (4x to 40x) magnification using light microscopy. However, the objective and microscope limitations only permitted imaging the superficial melanized nodules at 100x magnification, which provided a lower resolution of the nodules compared to the imaging of the extracted hemolymph. While in the case of *C. neoformans*, the nodules within the hemolymph appeared congruent to those viewed *in situ*, that might not always be the case. Nodules in extracted hemolymph during other fungal infection may not be entirely representative of those found throughout the entire larvae, so only viewing the hemolymph nodules may give a biased understanding of the fungal infection.

We also examined the melanization response to *Candida albicans* infection. *C. albicans* is known to trigger large scale systemic melanization in *G. mellonella* larvae ^33,44^. Similar to *C. neoformans*, we found melanized nodules in the hemolymph from larvae infected with *C. albicans*. Interestingly, the center of these nodules had melanized and smoothened areas that seemed more amorphous than those seen with *C. neoformans*, and additionally, we saw hyphal structures appeared less melanized than the spherical yeast-like structures. Using the tissue clarification method, we noted that the melanin-encapsulated *C. albicans* formed large rope-like aggregates without tissue tropism, with yeast being preferentially melanized over hyphal cells. Using *in vitro* time-lapse microscopy, we found that rapid melanization occurred, even in the absence of hemocytes. Additionally, after the melanization plateaus, the surviving fungus can break free from the melanin encapsulation and undergo melanin-evasive filamentation. This is followed by production of laterally-budding blastoconidium and a bloom in melanization around these newly formed yeast cells. Similar fungal morphologies and timelines were observed in dissected infected larvae, although the temporal kinetics were less resolved and identification of blastoconidium was less clear. Together, these data paint an interesting picture and allow insight into the pathogenesis of *C. albicans* within *G. mellonella* host. Hence, it appears that the melanin encapsulation can clear most of the yeast upon infection, however, cells that survive can then filament and evade subsequent melanin-mediated killing. The hyphae are known to penetrate and infect organs within the insect ^23^. The hyphae then produce yeast, which again triggers a burst of melanization that would likely cause damage to the surrounding tissue and eventually death of the organism.

In summary, we found evidence that the *G. mellonella* wax moth directly kills *C. neoformans* by encapsulating it with melanin *in vivo* using a GFP-expressing strains where fluorescence indicates viability. This association between melanin encapsulation and reduced viability provides the first direct evidence for fungal killing via melanin encapsulation *in vivo*. We also describe three different methodological approaches for studying the melanization response to fungi in *G. mellonella* and employ these techniques to study *C. neoformans* and *C. albicans* interactions with the melanin-based immune response. With *C. neoformans*, we show that both fungal melanins and fungal laccases can activate the insect’s melanization immune response, furthering our understanding of how these fungal components interact with insect immunity and alter the fungus’ pathogenesis. In *C. albicans*, we are able to observe how some melanin-encapsulated yeast are able to break through the melanization, and form melanin-evasive hyphae and pseudohyphae during infection. The direct association of insect melanization with antifungal defense further heighten concern that pesticides that inhibit the melanin reaction ^30^ could have untoward and unpredictable effects on insect populations.

## Acknowledgements

We would like to thank the entire Casadevall Lab for their contributions during lab meetings and other discussions of this project. We would like to thank Maryann Smith, Thomas Hitzelberger, and Kathy Spinnato for placing the years of weekly *G. mellonella* orders. Figure 6 was created using Biorender.com. D.F.Q.S., Q.D., and A.C. are funded by National Institute of Allergy and Infection Disease R01 AI052733. D.F.Q.S. is funded by National Institutes of Health 5T32GM008752-18 and 1T32AI138953-01A1. The funders had no role in study design, data collection and analysis, decision to publish, or preparation of the manuscript. The salaries of D.F.Q.S., Q.D., and A.C. are in part funded by the National Institute of Allergy and Infection Disease. The salaries of D.F.Q.S. and Q.D., are in part funded by the National Institutes of Health.

## Author contributions

Conceptualization – DFQS, QD, MK, JMH, AC; Methodology – DFQS, AC, QD, MK; Software – QD; Validation – DFQS; Formal analysis – DFQS, MK; Investigation – DFQS, MK; Resources – AC, JMH; Data curation – DFQS; Writing (Original Draft) – DFQS, AC; Writing (Review and Editing) – DFQS, QD, MK, JMH, AC; Visualization – DFQS, MK; Supervision – DFQS, JMH, AC; Project Administration – DFQS, AC; Funding Acquisition – JMH, AC

## Figure Legends

**Supplementary Figure 1.**
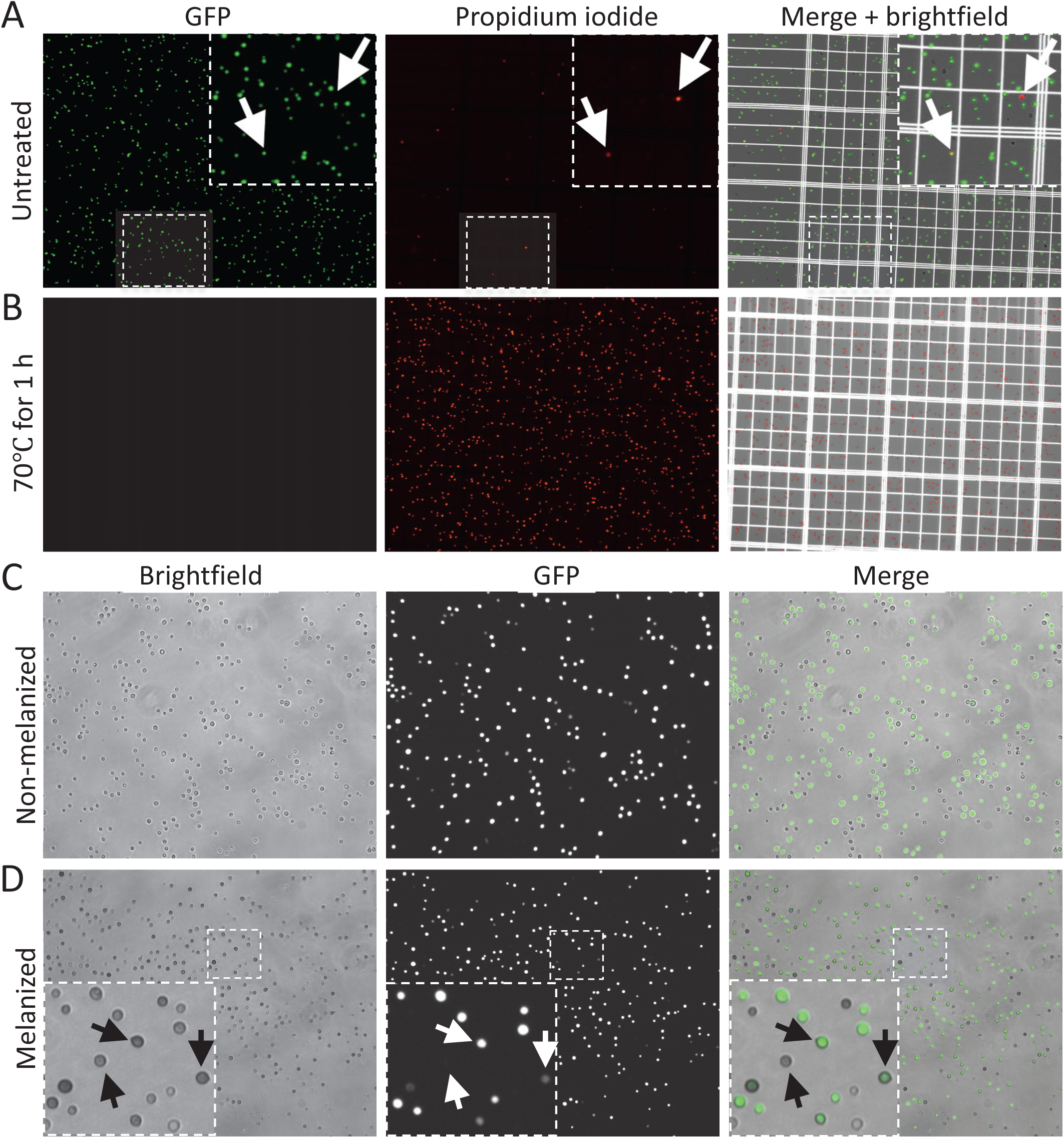
Fungal melanization does not affect GFP signal in the H99-GFP strain. (A-B) GFP signal is lost upon heat treatment for 1 hour, and cells become exclusively propidium iodine positive Insets in (A) indicate representative images of cells in untreated condition, with arrows pointing to a PI-only positive cell and a PI and GFP double positive cell. Non-melanized H99-GFP (C) and melanized H99-GFP (D) have comparable levels of GFP-fluorescence. Insets in (D) indicate representative selections of melanized cells that have GFP signal.

**Supplementary Figure 2.**
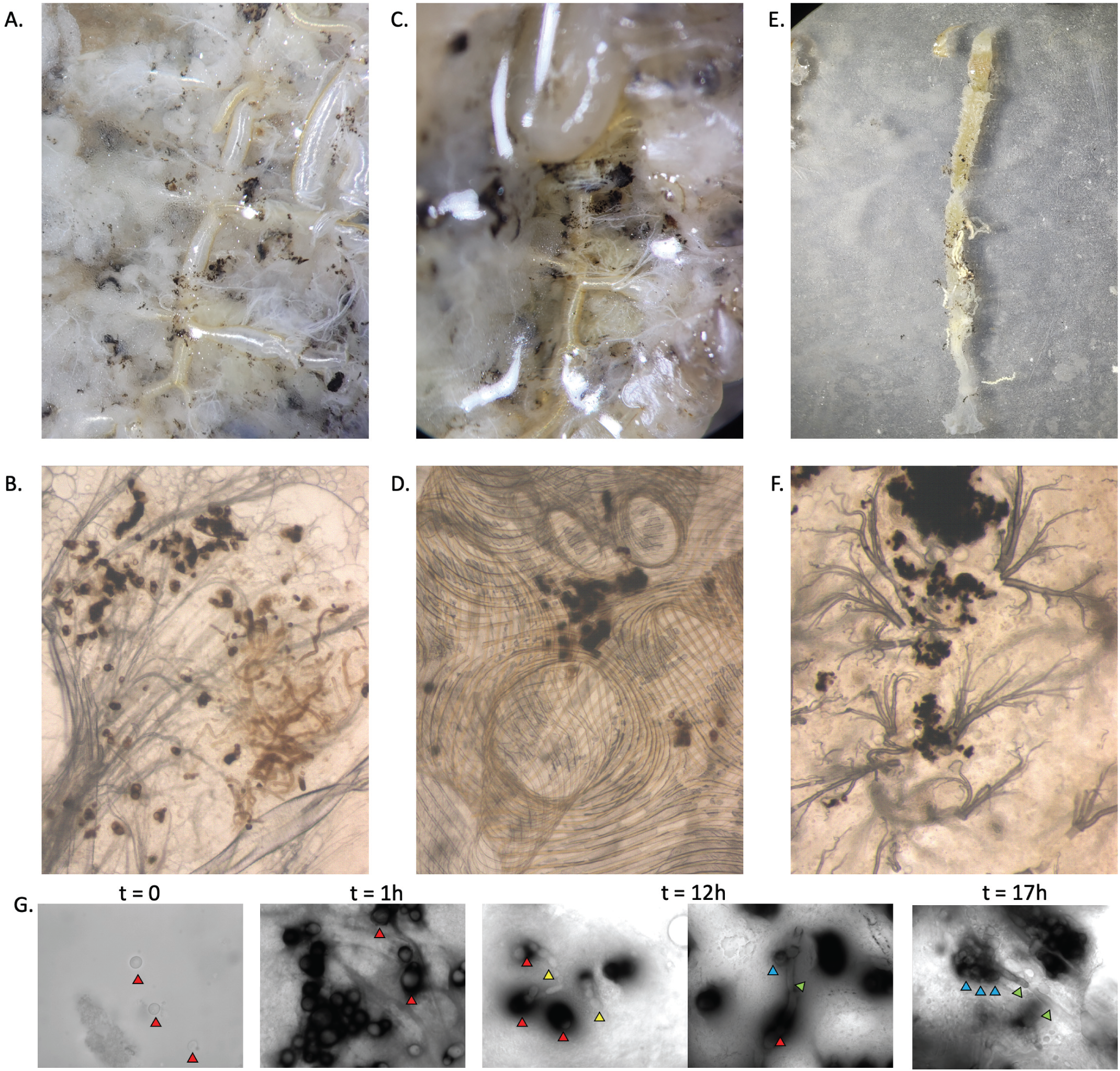
Dissection of *C. albicans* infected *G. mellonella*. There does not appear to be any specific tissue tropism for C. albicans infections in *G. mellonella*, with visible melanized nodules found in scattered in the Fat Bodies (A-C red arrow), trachea (A,C,D white arrow), and the gut (E,F). Microscopic analysis of the dissected tissue also reveals hyphal structures with reduced pigmentation compared to the yeast-like structures (B). (G) A similar progression of *C. albicans* morphological progression is seen in dissected tissues infections as is seen during the *in vitro* microscopy. At 0 minutes, there is no melanin encapsulation of the *C. albicans* yeast, whereas after 1 h, there is extensive melanization of *C. albicans* yeast. At 12 h, non-melanized hyphal and pseudohyphal structures are visible along with melanized yeast and potential laterally budding blastoconidium. By 17 h, there appears to be melanized laterally-budded blastoconidium with non-melanized hyphae, similar to what we see between hour 12 and 16 in the timelapse microscopy. Red arrows in (G) indicate yeast, yellow arrows indicate pseudohyphae, green arrows indicate hyphae, and blue arrows indicate potential laterally-budded blastoconidium.

**Supplementary Figure 3.**
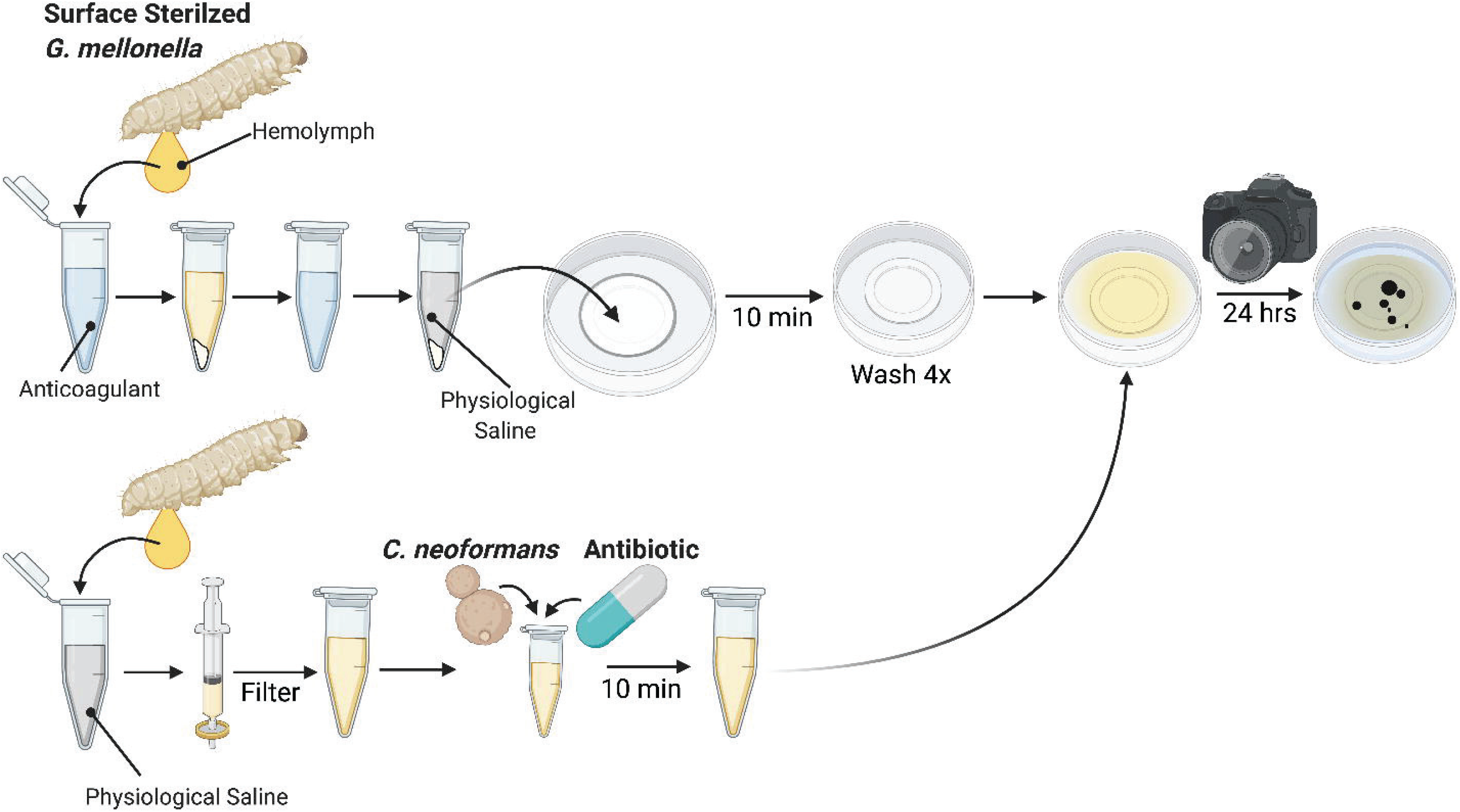
In vitro timelapse microscopy protocol. First, hemolymph from surface sterilized larvae are collected into anticoagulation buffer, hemocytes are centrifuged, washed in anticoagulant, and resuspended in insect physiological saline, and left to settle on a MatTek dish for 10 minutes. Simultaneously, hemolymph was collected into insect physiological saline, filtered with a PVDF 0.22 µm filter. Antibiotic and fungus is then added, left to sit for 10 minutes, and added to the adherent hemocytes following 4 washes with insect physiological saline. Timelapse microscopy was then performed for up to 24 hours.

## Materials and Methods

### Biological materials

*G. mellonella* last-instar larvae were obtained through Vanderhorst Wholesale, St. Marys, Ohio, USA. *C. neoformans* strain H99 (serotype A), *C. neoformans* strain H99-GFP ^45^, *C. neoformans lac1Δ* mutant, and *Candida albicans* strain 90028 were kept frozen in 20% glycerol stocks and sub-cultured into yeast peptone dextrose (YPD) broth for 48 h at 30°C prior to each experiment. For H99-GFP infections, frozen stock was streaked out first onto YPD agar, and green colonies were inoculated into YPD broth for 48 h at 30°C prior to each experiment. The yeast cells were washed twice with PBS, counted using a hemocytometer (Corning, New York, USA), and adjusted to 10^7^ cells/ml for an injection inoculum of 1 × 10^5^ cells/larva. *C. albicans* infections were performed at 5 × 10^5^ cells/larva.

### Extraction of hemolymph from fungal*-*infected *Galleria mellonella* larvae

Infection of *G. mellonella* larvae was performed as previously described ^30^. Briefly, washed *C. neoformans* or *C. albicans* cultures, resuspended to 10^7^ cells/ml were injected in the right rear proleg of larvae ranging from 175 to 225 mg. Infected larvae were then incubated at 30°C. Three days following infection, larvae were removed from incubator, and hemolymph was extracted by puncturing the right rear proleg with an 18 G needle. Removed hemolymph from 3 larvae was collected directly into 1 ml anticoagulation buffer at room temperature ^46^. Hemolymph was centrifuged for 5 minutes at 4,000 x *g* and resuspended in 200 µl insect physiological saline (IPS) (150 mM sodium chloride, 5 mM potassium chloride, 7.21 mM calcium chloride, 1 mM sodium bicarbonate, pH 6.90 – adapted from ^47–49^). Samples were placed on slides and nodules were imaged using Olympus AX70 microscope with a 100x oil immersion objective.

### *C. neoformans* GFP Viability Assay

H99-GFP strain was streaked from frozen stock on YPD agar and incubated at 30°C. 2 ml YPD was inoculated with H99-GFP and incubated for ∼18 h at 30°C with rotation. Culture was diluted to OD 0.5 and 100 µL was incubated at 70°C for 1 h using a thermocycler. 100 µL of untreated and heated samples were stained with 10 µg/ml propidium iodide (Invitrogen). 10 μL of stained samples were loaded onto a hemocytometer and imaged using a 10X objective and Zeiss AxioImager M2 (60x Olympus objective) equipped with a Hamamatsu Orca R2 camera and Volocity Software (Perkin Elmer). Images were analyzed using ImageJ/FIJI software. Fluorescence channel images were processed by adjusting the minimum pixel value to 10 and maximum to 90. Number of fluorescent cells for each channel were counted using Measure Particles. Number of double fluorescence positive and double fluorescence negative cells were enumerated manually.

### *C. neoformans* GFP Fungal Survival Assay *in vivo*

*G. mellonella* larvae were infected as previously described using H99-GFP. Larvae were incubated for 3 days at 30°C, and hemolymph was extracted. Melanized nodules were visualized using an Olympus AX70 microscope with 488 excitation/520 nm emission fluorescence microscopy to visualize the GFP signal. Images were taken at 100x magnification with the same exposure, and manually marked as positive or negative for GFP fluorescence, and melanin-encapsulated or unencapsulated. For fluorescence and melanin intensity measurements, images were analyzed using the Measure tool in FIJI ^50^, and the 8-bit mean gray value of each cell was measured in both channels. The region selected for the melanin measurements extended the edge of the fungal capsule, and the GFP intensity measurements were from selections limited to the fluorescent cell’s body.

### Phenoloxidase Inhibition and Fungal Survival Assay *in vitro*

Serial dilutions of phenylthiourea (PTU) were performed in 100 µl IPS buffer, to which 5 µl of 10^6^ cells of *C. neoformans* was added. *G. mellonella* hemolymph was extracted as previously described into insect physiological saline, and 100 µl of the mixture was added to each well. The mixture was incubated at room temperature for 24 hours protected from light. Following the incubation, the contents of each well were resuspended and diluted 1:16 in PBS. From the dilution, 5 µl was spotted on a Sabouroud agar plate. The plate was incubated at 30°C for 24 h and colonies were enumerated under a dissection microscope.

### Tissue Clearing of *Galleria mellonella* following fungal infection

*G. mellonella* larvae were infected with *C. neoformans* or *C. albicans* as described above. Five days following infection, groups of three larvae were removed from incubator and injected with 10 µl of 1 M ascorbic acid to inhibit new melanization and oxidation of endogenous catecholamines during the tissue clearing process. Ten minutes following the ascorbic acid injection, larvae were placed at -20°C for fifteen minutes to euthanize them, then injected with an additional 10 µl of 1 M ascorbic acid. Larvae were immediately placed in 40 mL of 4% paraformaldehyde. Larvae were fixed, permeabilized, and cleared in Benzyl Alcohol and Benzyl Benzoate (BABB) solution as previously described ^32^. Following 5 to 7 days of tissue clearing, larvae were removed from the BABB solution and pressed between two glass microscope slides. Once flattened, a coverslip was placed on top of the larvae and parafilmed into place. Larvae were imaged using Olympus AX70 microscope with 4x, 20x, and 100x objectives.

### Imaging *Galleria mellonella* hemocytes *in vitro*

To collect and isolate hemocytes *G. mellonella* larvae were surface sterilized in two sequential baths of 70% ethanol, followed by 10% bleach, then dried on sterile paper towels. Five to 10 drops of hemolymph were extracted as described above into room temperature anticoagulation buffer and inverted 3 times. Hemolymph was centrifuged at 400 x *g* for 4 minutes, the supernatant was removed, and the hemocytes were resuspended in 1 ml anticoagulation buffer and centrifuged. The supernatant was completely removed and hemocytes were resuspended in 200 µl of insect physiological saline (IPS). The 200 µl suspension of hemocytes were added to the coverslip of a MatTek dish and allowed to settle for 10 minutes. Following the 10 minutes, the buffer and unsettled hemocytes were removed, and the coverslip was washed 4 times with 1 ml of IPS. The hemocytes are seeded into the coverslip at a cell density of 1.5 × 10^6^ cells/ml and the resulting hemocyte density after washing is approximately 2-3 × 10^3^ cells/mm^2^.

While the hemocytes were being isolated, cell-free hemolymph was being prepared. Approximately 10 drops of hemolymph were removed from *G. mellonella* larvae and collected directly into 1 ml IPS. To remove hemocytes, the mixture was filtered using a 0.22 µm syringe-driven PVDF filter. Cell-free hemolymph was stored up to a week at -80C. Penicillin-Streptomycin (Gibco, Thermo Fisher) antibiotic was added at 1x concentration to the cell-free hemolymph. For experiments looking at the interaction of hemocytes with fungi or a virulence factor, the cells or component are added at this stage.

Following the hemocyte washes, 1 ml of cell-free hemolymph was added to the entirety of the MatTek dish, followed by an addition 1 ml of IPS. The MatTek dish was covered and imaged using the OpenFlexure microscope and software and time-lapse microscopy was performed every minute for 16-24 hours ^51^. This protocol is summarized in Supplementary Figure 3.

All timelapse data was analyzing using FIJI ^50^ and particle measurements were made by converting the image sequence to 8-bit, setting a threshold of 0-50 gray value, and analyzing any particle over the size of 4 pixels^2^. Measuring the time until germ tube formation was done manually by recording the frame in which the first of the germ tube was visible.

### Melanin Ghost Isolation

*C. neoformans* cultures were grown in minimal media with 1 mM L-DOPA for 7 days at 30°C. Cells were collected and mixed 1:1 with 12 N hydrochloric acid (HCl), for a final concentration of 6 N HCl. Cells were heated for 1 hour at 85°C under constant shaking at 350 RPM. Control cells were heat killed cells were incubated for 1 hour at 85°C in PBS. Cells were washed twice in PBS and subsequently used in the time-lapse microscopy.

## Multimedia Files

Supplementary Video 1_C. neoformans timelapse

Supplementary Video 2_C. neoformans timelapse

Supplementary Video 3_Melanin Ghost vs heat killed

Supplementary Video 4 _Melanin ghost without hemocytes

Supplementary Video 5_Melanin ghost timelapse

Supplementary Video 6_Hemocyte-ghost interactions

Supplementary Video 7_In situ nodule projection

Supplementary Video 8_Melanin Bloom Candida

Supplementary Video 9_Candida albicans escape

Supplementary Video 10_C. neoformans Anticoagulation Buffer

Supplementary Video 11_No fungus timelapse

